# Auditory deviance detection in the human insula: An intracranial EEG study

**DOI:** 10.1101/487306

**Authors:** Alejandro O. Blenkmann, Santiago Collavini, James Lubell, Anaïs Llorens, Ingrid Funderud, Jugoslav Ivanovic, Pål G. Larsson, Torstein R. Meling, Tristan Bekinschtein, Silvia Kochen, Tor Endestad, Robert T. Knight, Anne-Kristin Solbakk

## Abstract

While the human insula is known to be involved in auditory processing, knowledge about its precise functional role and the underlying electrophysiology is limited. To assess its role in automatic auditory deviance detection we analyzed the high frequency EEG activity (75-145 Hz) from 90 intracranial insular electrodes across 16 patients who were candidates for resective epilepsy surgery while they passively listened to a stream of standard and deviant tones. Deviant and standard tones differed in four physical dimensions: intensity, frequency, location and time. Auditory responses were found in the short and long gyri, and the anterior, superior, and inferior segments of the circular sulcus of the insular cortex, but only a subset of electrodes in the inferior segment showed deviance detection responses, i.e. a greater and later response to deviants relative to standards. Altogether, our results indicate that the human insula is engaged during auditory deviance detection.

## INTRODUCTION

The role of the human insula in auditory processing is incompletely understood. Evidence from intracranial EEG (iEEG) indicates that the insula plays a role in auditory processing and that when stimulated electrically it elicits auditory illusions and hallucinations (Afif, Minotti, Kahane, & Hoffmann, 2010; Zhang et al., 2018). Previous studies reported auditory agnosia after bilateral insular lesion (Bamiou, Musiek, & Luxon, 2003) and deficits in the temporal resolution and sequencing of sounds after unilateral insular stroke (Bamiou et al., 2006). Additionally, insular neurons fire in response to auditory stimuli in non-human primates (Bieser, 1998; Remedios, Logothetis, & Kayser, 2009), which share a similar auditory cortex architectonics with humans (Fullerton & Pandya, 2007; Galaburda & Sanides, 1980).

Responses to unexpected sounds in regular auditory streams have been studied with the mismatch negativity (MMN), an event-related potential (ERP) that peaks around 150-250 ms after the onset of an infrequent acoustic stimulus (Näätänen, Paavilainen, Rinne, & Alho, 2007). MMN is considered a prediction error signal reflecting automatic detection (Bubic, 2010; Clark, 2013). Multiple iEEG studies have shown that the posterior part of the superior temporal plane is involved in deviance detection (Edwards, Soltani, Deouell, Berger, & Knight, 2005; El Karoui et al., 2015; Halgren et al., 1995; Phillips, Blenkmann, Hughes, Kochen, Bekinschtein, Cam-CAN, et al., 2016), as well as the lateral prefrontal cortex and the nucleus accumbens (Durschmid et al., 2016a, 2016b).

Previous fMRI (Nazimek, Hunter, Hoskin, Wilkinson, & Woodruff, 2013; Sabri, Kareken, Dzemidzic, Lowe, & Melara, 2004; Schall, Johnston, Todd, Ward, & Michie, 2003), PET (Müller, Jüptner, Jentzen, & Müller, 2002), and MEG studies (Lappe, Steinsträter, & Pantev, 2013) have argued that the human insula plays a role in automatic deviance detection. However, a high spatial and temporal resolution description of its electrophysiological activity is lacking. In the current study, we addressed this issue by analyzing High Frequency Activity (HFA; 75-145 Hz) in iEEG recordings, a reliable electrophysiological correlate of underlying averaged spiking activity generated by the thousands of neurons that are in the immediate vicinity of the recording electrodes (Lachaux, Axmacher, Mormann, Halgren, & Crone, 2012; Ray & Maunsell, 2011, Rich & Wallis, 2017; Watson, Ding, & Buzsáki, 2018).

## METHODS

### Ethics statement

This study was approved by the Research Ethics Committee of El Cruce Hospital, Argentina, and the Regional Committees for Medical and Health Research Ethics, Region North Norway. Patients gave informed written consent prior to participation.

### Participants

We studied 16 normal hearing adults (6 female, mean age = 31, range 19-50 years) with drug-resistant epilepsy who were potential candidates for resective surgery. Patients underwent invasive stereo electroencephalography (SEEG) recordings as part of their pre-surgical evaluation. Intracranial depth electrodes were temporarily implanted to localize the epileptogenic zone and eloquent cortex. Data were collected at El Cruce Hospital or Oslo University Hospital. Patients were implanted with depth electrodes of 8-18 contacts with 1.5-5 mm inter-electrode distance (AdTech, USA, and DIXI Medical, France). We selected datasets with insula coverage (median 4.5, range 1-21 contacts per patient).

### Task

A “Optimum-1” multi-dimensional auditory oddball paradigm was used (Näätänen, Pakarinen, Rinne, & Takegata, 2004; Phillips et al., 2016). Standard tones were defined across four dimensions: frequency, intensity, location and time. Standards were interleaved with deviant tones that deviated in one of the four dimensions, while holding other stimulus dimensions constant. Table 1 depicts the most relevant stimulus features for all conditions. Tones were presented every 500 ms, in blocks of 5 minutes consisting of 300 standard and 300 deviant tones. Fifteen standards were played at the beginning of each block. Deviants were presented in pseudo-random order such that a deviant type never appeared twice in a row (Figure 1A). Participants were asked not to pay attention to the sounds while reading a book or magazine. They completed 3 to 10 blocks, providing at least 1800 trials. The tones were presented through headphones using Psychtoolbox-3 (Kleiner, Brainard, 2007) for Matlab (The MathWorks Inc., USA).

**Figure 1.**
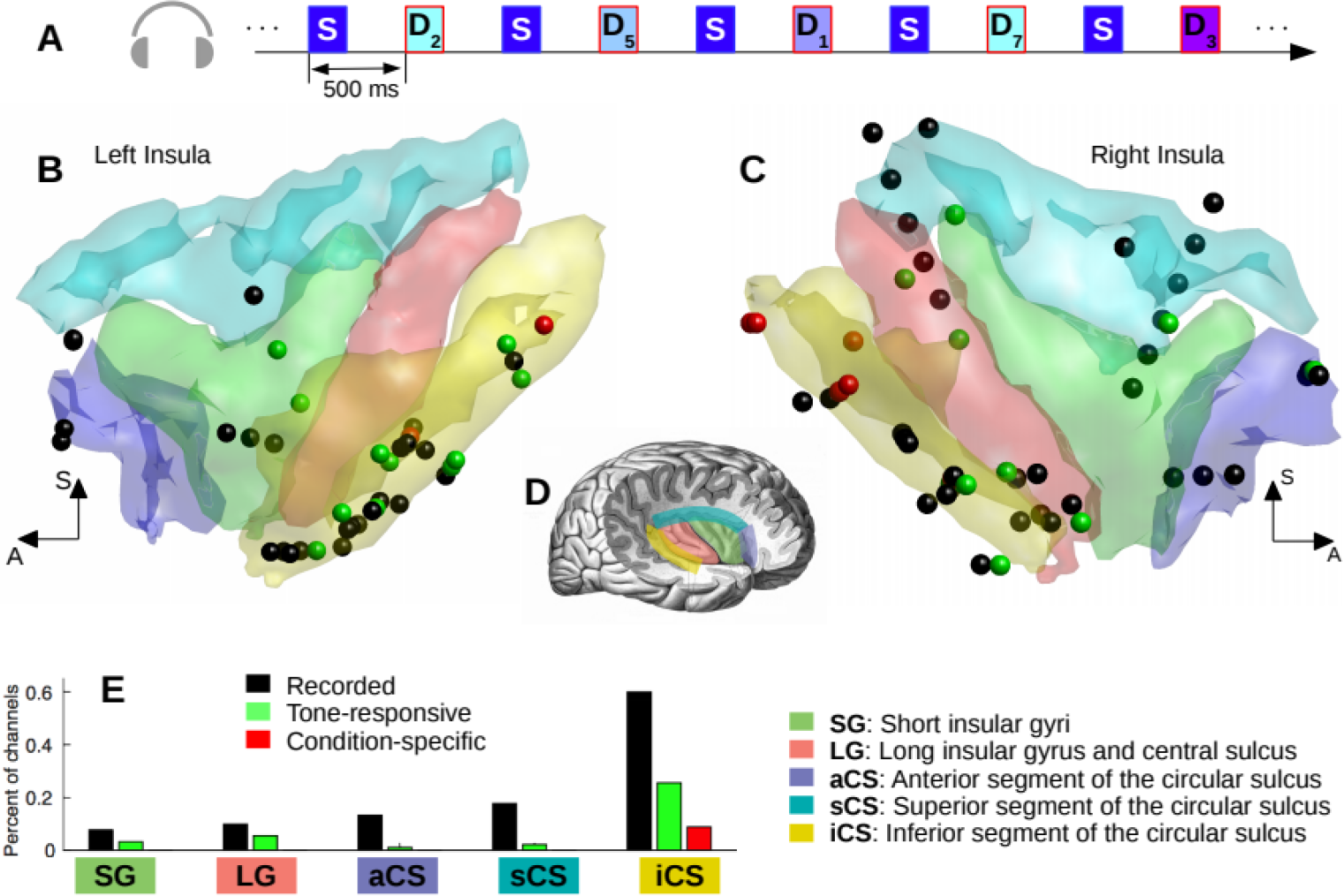
Experimental design and spatial distribution of recorded, responsive and task-condition specific channels. **A.** Experimental design denoting standard and deviant tones are interleaved. Deviants in different physical dimensions (frequency, intensity, location and time) were randomly distributed. **B & C.** Recorded (black), tone responsive (green), and task-condition specific (red) channels from all subjects are shown over a 3D reconstruction of the left and right insular cortex. The anatomical reconstruction with subdivisions was obtained from an MRI segmentation of the brain scan of one patient (OSL14). **D.** Illustrative map of the insular subdivisions. Modified from Textbook and Atlas of Human Anatomy (Sobotta, 1908). **E.** Anatomical distribution of recorded, responsive and condition specific channels by anatomical area.

**Table 1.** Stimulus features. Empty cells indicate same features as standard tones.

| Mismatch Dimension | Condition | Frequencies | Intensity | Sound Location | Time |
| --- | --- | --- | --- | --- | --- |
|  | Standard | 500, 1000, 1500 Hz | 0 dB | Frontal | Duration 75 ms |
| Frequency | Freq-High<br>Freq-Low | 550, 1100, 1650 Hz<br>450, 900, 1350 Hz |  |  |  |
| Intensity | Int-Up<br>Int-Down |  | +10 dB<br>-10 dB |  |  |
| Location | Loc-Left<br>Loc-Right |  |  | Left<br>Right |  |
| Time | Time-Dur<br>Time-Gap |  |  |  | Duration 25ms<br>7ms gap |

### Data acquisition

Pre-implantation structural MRI and post-implantation CT scans were acquired for each participant. SEEG data were recorded using an Elite (Blackrock NeuroMed LLC, USA), a NicoletOne (Nicolet, Natus Neurology Inc., USA), or an ATLAS (Neuralynx, USA) system with sampling frequencies of 2000, 512, and 16000 Hz respectively.

### Electrode localization

Post-implantation CT images were coregistered to pre-implantation MRI images using SPM12 (Studholme, Hill, & Hawkes, 1999). MRI images were processed using the FreeSurfer standard pipeline (Dale, Fischl, & Sereno, 1999), and individual cortical parcellation images were obtained using the Destrieux atlas (Destrieux, Fischl, Dale, & Halgren, 2010). Images were spatially normalized to the MNI-152 template using SPM12 (Ashburner & Friston, 2005) and electrode coordinates were obtained using the iElectrodes toolbox (Blenkmann et al., 2017). Anatomical labels were automatically assigned to each contact based on the Destrieux atlas using the aforementioned toolboxes, and confirmed by a neurologist (RTK).

### Signal pre-processing

Raw iEEG recordings were manually inspected and channels or epochs showing epileptiform activity or abnormal signal were removed. Channels located in lesional tissue or tissue that was later resected were also excluded. Bipolar channels were computed as the difference between signals recorded from pairs of neighboring electrodes in the same electrode array. Subsequently, we refer to the bipolar channels as “channels”. Data were low-pass filtered at 180 Hz and line noise was removed using bandstop filters at 50, 100, and 150 Hz. Data were then segemented into 2000 ms epochs (-750 ms before and 1250 ms after tone onset) and demeaned. We rejected trials with a raw amplitude above five standard deviations from the mean for more than 25 consecutive ms, or with a power spectral density above five standard deviations from the mean for more than 6 consecutive Hz. Data were resampled to 1000 Hz.

### Region of interest channel selection

We studied channels in the insular cortex categorized into the following sub-areas based on Duvernoy’s anatomical nomenclature (Destrieux et al., 2010; Duvernoy, 1999; Figure 1A-C: i) short insular gyri (SG), ii) long insular gyrus and central insular sulcus (LG), iii) anterior segment of the circular sulcus (aCS), iv) superior segment of the circular sulcus (sCS), and v) inferior segment of the circular sulcus (iCS). Ninety insular electrodes were analyzed across 16 patients. The distribution of channels across sub-areas is shown in Fig 1E and across patients in Supplementary Table 1.

### High frequency activity extraction

Preprocessed data were filtered into eight bands of 10Hz bandwidth ranging from 75 to 145Hz using bandpass filters. The Hilbert transform was applied to each filtered signal to create a complex-value analytic time series. The modulus of these signals was the analytic amplitude time series corresponding to specific frequency bands. The mean amplitude of the baseline period (-100 to 0 ms) of each trial and frequency band was removed from the entire trial. The mean of the eight frequency bands was computed such that one time series was built per trial. Finally, for each channel, all trial time series were divided by the standard deviation pulled from all trials in the baseline period, thereby computing the HFA time series of each channel as a normalized measure that was relative to the baseline activity.

#### Tone-responsive channels

Channels were considered responsive if the median HFA response to tones, irrespective of type, was statistically higher in the post-stimulus period (0 to 300 ms after stimulus onset) relative to the baseline period (-100 to 0 ms) (Durschmid et al., 2016a). We tested the differences in five 100 ms overlapping windows in the post-stimulus period, with 50% overlap, using a permutation based approach. For each channel, we computed the difference between the HFA medians in the post-stimulus windows and the HFA medians in the baseline period. We then created an empirical distribution by circular-shifting of the HFA trial time series (between -100 and 500 ms) by a random number of samples. This method allows any time-locked neural activity to be teased apart. Differences between the medians in the post-stimulus and baseline periods were measured for each surrogate set of trials. The procedure was repeated 1000 times and a null distribution of differences was built. Channels exceeding the 97.5th percentile of the channel-specific surrogate distribution in any of the five windows were considered showing a significant time-locked HFA modulation and were marked as Tone-responsive channels.

#### Condition-specific channels

To determine whether the Tone-responsive channels were sensitive to intensity, frequency, location or time deviances, we performed for each channel a one-way ANOVA cluster-based permutation test with 3 levels: the standard tones and the two deviant tones in each dimension (alpha = 0.05, 1000 permutations). Post-hoc one-tailed t-tests were performed to determine if the HFA responses to deviants were higher in amplitude than those to standards (cluster-based permutation test, alpha = 0.025, 1000 permutations).

Pre-processing and statistical analysis was performed in Matlab using the Fieldtrip toolbox (Oostenveld, Fries, Maris, & Schoffelen, 2011) and custom code. All filters used had zero phase distortion.

## RESULTS

All participants reported that they were able to focus on the provided reading material and did not attend to the tones during the experiment. When queried, participants reported that they did not notice any pattern in the stream of tones.

### Auditory effects

Twenty-nine channels responded to tones in 11 out of the 16 patients (mean 2.6, range 1-5 channels per patient, permutation test, p < 0.025, Figure 1B-C). Channels that showed a significant response were non-uniformly distributed in all subdivisions of the insular cortex; the predominant number of channels were located in the iCS (Figure 1E; Fisher’s exact test, p = 0.02). Figure 2A shows the mean HFA response of one illustrative electrode, along with the single trial HFA and corresponding significant period. In Figure 2C the mean response of the Tone-responsive channels averaged across tone types (n = 29) can be seen. Median peak response latency was 120 ms, 95% CI [100, 141] ms.

**Figure 2.**
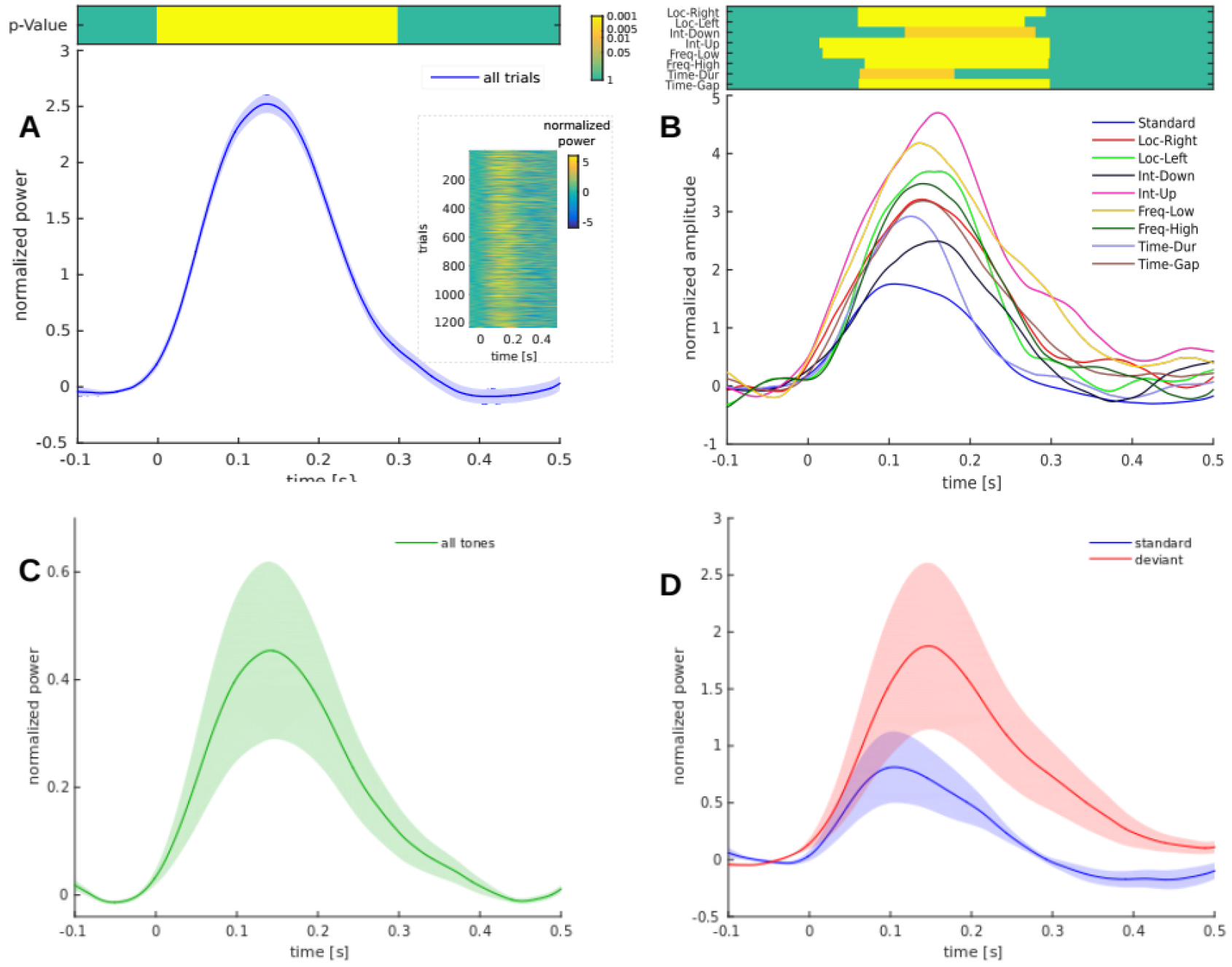
HFA responses. **A.** Mean HFA response to all tones in one illustrative Tone-responsive channel (see Methods, section *Tone-responsive channels* for definition). Inset shows the single trials time courses. Top image shows the active period HFA with significantly higher amplitude than baseline (p-values obtained from permutation test). **B.** Mean HFA response to each condition in one Condition-specific channel. The top image denotes the periods where deviant HFA responses were significantly higher than the standard tone responses (post hoc permutation test). **C.** Average HFA response to tones obtained from all Tone-responsive channels. **D.** Average HFA response to standard (blue) and deviant (red) tones from all Condition-specific channels (see Methods, section *Condition-specific channels* for definition). Shadowed areas depict mean ± SEM.

To obtain a global description of the role of the insula in deviance detection we tested the HFA response to standards versus deviants in all Tone-responsive channels. Deviants showed a higher response in the 134-196 ms interval after tone onset (permutation test, p = 0.014, Supplementary Figure 1).

### Deviance detection effects

Eight channels across five patients showed condition-specific effects to one or more of the deviant dimensions (permutation test, p < 0.05). All Condition-specific channels were located in the iCS (Figures 1B-E), which represent 15% of the recorded, and 32% of the Tone-responsive channels in the area. Figure 2B shows the HFA responses to all stimulus types in one illustrative channel.

Across the eight channels, seven had significantly higher responses to at least one deviant condition compared to standards (post-hoc permutation test). For illustration, we computed the mean HFA time course (averaged across significant channels) of each deviant condition and its corresponding standard tone response. *Loc-Left* (3 channels, p < 0.001) and *Loc-Right* (3 channels, p < 0.001) deviant responses showed similar time courses independent of tone laterality (Figure 3A). The time course responses for *Freq-High* (5 channels, p < 0.001) and *Freq-Low* (4 channels, p < 0.05) deviants were also similar (Figure 3C). However, the *Int-Up* (5 channels, p < 0.001) deviant produced a larger response than the *Int-Down* (3 channels, p < 0.05) deviant, but the latter elicited a larger response than standards, suggesting that the HFA response was not solely driven by tone intensity (Figure 3B). *Time-Gap* (5 channels, p < 0.001) and *Time-Dur* (4 channels, p < 0.05) deviants yielded different HFA time courses (Figure 3D). Supplementary Figures 2-5 show the spatial distribution of these effects. Additionally, one channel showed a higher HFA response to standards when compared to *Int-Down* deviants. Condition specific responses were concurrent in some channels. Three of the channels were sensitive to all stimuli dimensions, two channels to two dimensions, and two channels to only one dimension.

**Figure 3.**
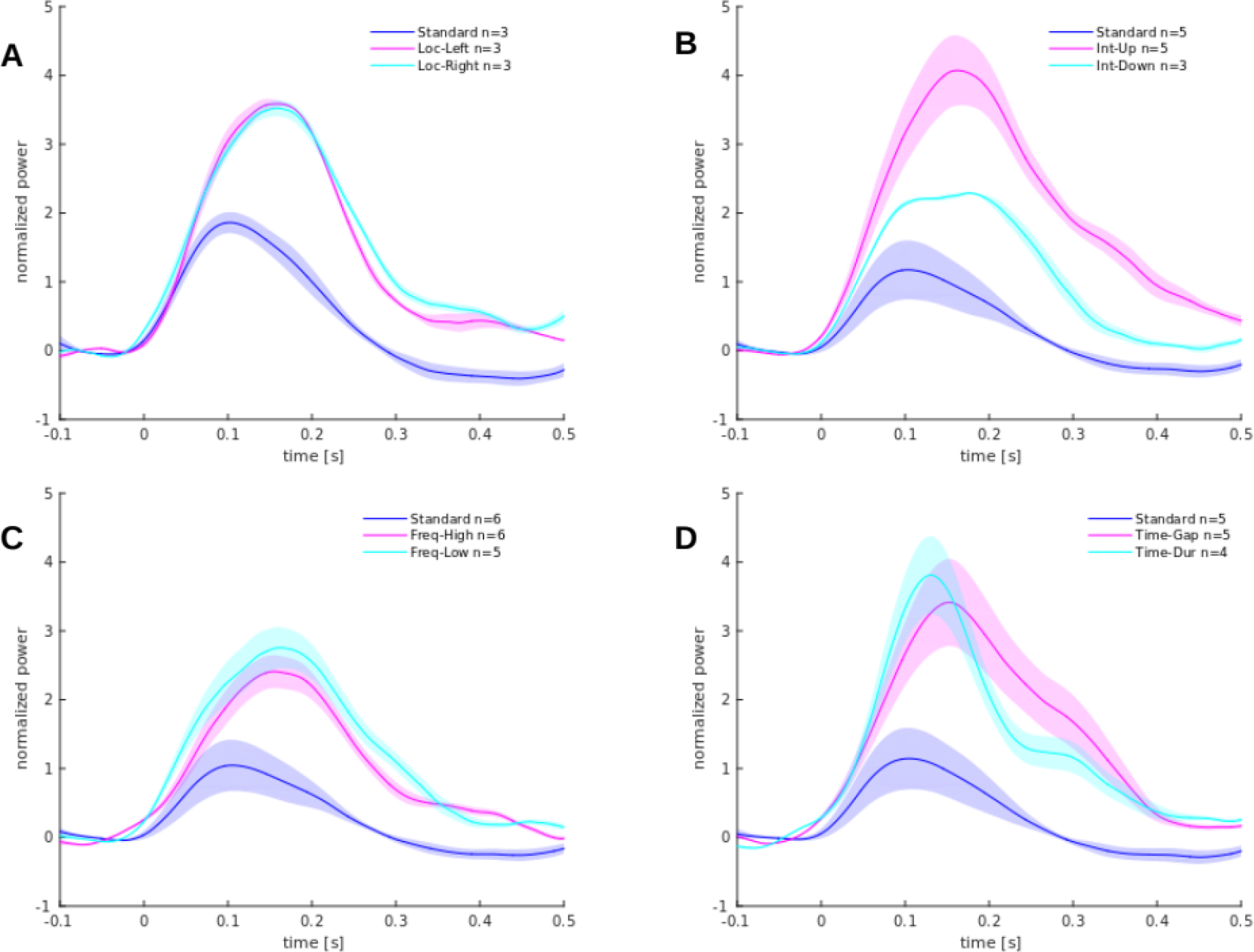
Condition specific HFA responses in the inferior segment of the circular sulcus of the insula. **A.** HFA responses to Loc-Left (magenta) and Loc-Right (cyan) deviant tones, and standard tones (blue). **B.** HFA responses to Int-Up (magenta) and Int-Down (cyan) deviant tones, and standard tones (blue). **C.** HFA responses to Freq-High (magenta) and Freq-Low (cyan) deviant tones, and standard tones (blue). **D.** HFA responses to Time-Gap (magenta) and Time-Dur (cyan) deviant tones, and standard tones (blue). *n* indicates the number averaged of channels. Shadowed areas depict mean ± SEM.

We computed the mean HFA response of Condition-specific channels to standards and deviants (all types merged) (Figure 2D). Standard tones showed an earlier peak latency than deviants (median 105 vs. 143 ms, Wilcoxon signed rank test, p = 0.008, Supplementary Figure 6).

To exploit the high spatial resolution of iEEG, we divided the iCS into three parts along the anterior-posterior axis and analyzed the spatial distribution. Deviance detection channels were located in the posterior and middle parts, but not the anterior part. See supplementary material for details.

## DISCUSSION

We examined iEEG responses in the human insula to standard and deviant tones. The results revealed that HFA tracked both auditory processing and automatic deviance detection in the insular cortex.

### The insula as part of the auditory network

Overall, our findings support prior evidence in showing that the insular cortex is involved in auditory processing and indicates that predominantly the iCS and LG areas of the insula are responsible. Previous studies have shown that posterior insula auditory responses resemble those observed in Heschl’s gyrus (Zhang et al., 2018), and that auditory perception is altered by electrical stimulation (Afif et al., 2010; Zhang et al., 2018) or focal insular strokes (Bamiou et al., 2003; Habib et al., 1995). Single unit responses in non-human primate insular cortex have been shown to encode amplitude and frequency modulated sounds (Bieser, 1998), and vocal communication sounds in its posterior part (Remedios et al., 2009). Additionally, the posterior insula is strongly connected to the primary auditory cortices (Ghaziri et al., 2017; Uddin, Nomi, Hébert-Seropian, Ghaziri, & Boucher, 2017; Zhang et al., 2018), and its cytoarchitectonics is similar to that of cortical sensory areas (Rivier & Clarke, 1997).

### The role of the insula in automatic auditory deviance detection

Prior iEEG studies have suggested that the posterior part of the superior temporal plane is a key contributor to the scalp MMN observed during automatic deviance detection (El Karoui et al., 2015; Halgren et al., 1995, Edwards et al., 2005; Kropotov et al., 1995; Rosburg et al., 2005). Additionally, results from fMRI (Opitz, Rinne, Mecklinger, von Cramon, & Schröger, 2002) and MEG/EEG source localization studies (Lappe et al., 2013; Rinne, Alho, Ilmoniemi, Virtanen, & Näätänen, 2000) have also identified the posterior part of the superior temporal plane as a source for the MMN. Furthermore, other studies have implicated the frontal cortex in deviance detection (Bekinschtein et al., 2009; Liasis, Towell, Alho, & Boyd, 2001; Phillips et al., 2016; Rosburg et al., 2005; Durschmid et al., 2106a, 2016b, Deouell. 2007). As of yet, no iEEG study has reported data from the human insula during deviance detection.

In our study, the results indicate that the iCS is involved in automatic auditory deviance detection. The HFA responses of the iCS were sensitive to deviations in the frequency, intensity, location, and time dimensions. In agreement with our findings, both fMRI and lesion studies have suggested involvement of the insular cortex in deviance detection. Griffiths et al. (1996) reported that a right insular stroke patient was unable to detect the auditory source of movements, and Bamiou et al. (2006) reported that unilateral insular stroke patients had deficits in temporal resolution and sequencing of sounds. From fMRI studies it has been shown that the left iCS responds to non-attended sound deviants in the frequency domain (Sabri et al., 2004), that the left anterior insula responds to tone duration deviants (Schall et al., 2003), and that there is a greater activation in the left insula for unexpected versus expected attended sounds (Nazimek et al., 2013). However, due to limited spatial resolution, fMRI and lesion studies cannot rule out the possibility that the temporal cortices adjacent to the insula are contributing to the activations (fMRI) or functional deficits (lesion). In contrast, intracranial HFA obtained from a bipolar montage provides the spatial and temporal resolution necessary to localize the source and timing of local neural activity (Lachaux et al., 2012; Ray & Maunsell, 2011; Rich & Wallis, 2107; Watson et al., 2018). Hence, our study specifically identified the iCS as the sub-insular area involved in deviance detection, showed its sensitivity to auditory deviance in multiple physical dimensions, and its temporal response profile.

Extending our results to other findings, HFA responses to standard and deviant tones differed from each other at a latency that is concurrent with the typical MMN’s peak latency (Garrido et al., 2009; Näätänen et al., 2007), revealing a common ground for the ERP and HFA analysis. Interestingly, the different peak latencies for deviants and standards indicate that a more elaborate process is unfolding in situations where sensory predictions are violated. In the same vein, higher responses to *Int-Down* deviants when compared to standards are difficult to explain under the adaptation hypothesis (i.e. that responses are due to fresh-afferent neurons) and strongly suggest a predictive coding process.

## Supporting information

## ACKNOWLEDGMENTS

We appreciate the patients for kindly participating in our study. We thank Julia Kam and other members of the Knight Lab for valuable discussions. We want to express our gratitude to the EEG technicians at El Cruce Hospital and Rikshospitalet for their support. This work was partly supported by the Research Council of Norway project number 240389.

