## Supplementary material for "Auditory deviance detection in the human insula: An intracranial EEG study"

**Demographic characteristics of patients with implanted electrodes**

| Pa-tient | Sex | Age | Insular channels | | | | | Total channels |
|---|---|---|---|---|---|---|---|---|
| | | | **SG** | **LG** | **aCS** | **sCS** | **iCS** | |
| 1 | M | 20 | | | | | 2 | 2 |
| 2 | F | 22 | 1 | 3 | | 1 | 4 | 6 |
| 3 | M | 19 | | | | | 3 | 3 |
| 4 | F | 49 | | 2 | | | 6 | 8 |
| 5 | M | 19 | | | 2 | | 3 | 5 |
| 6 | M | 21 | 3 | | | | 5 | 8 |
| 7 | M | 25 | | | | | 2 | 2 |
| 8 | M | 37 | | | | | 5 | 5 |
| 9 | F | 23 | | | 3 | | | 3 |
| 10 | M | 38 | 3 | 1 | | 2 | | 3 |
| 11 | F | 30 | | | 2 | | 8 | 10 |
| 12 | M | 27 | | | | | 3 | 3 |
| 13 | F | 34 | | | | | 6 | 6 |
| 14 | M | 38 | | 3 | 5 | 11 | 4 | 21 |
| 15 | M | 48 | | | | 1 | | 1 |
| 16 | F | 50 | | | | 1 | 3 | 4 |
| **Total** | 6F/10M | 31.3 | **7** | **9** | **12** | **16** | **54** | **90** |

**Supplementary table 1.** Demographic characteristics of implanted patients and number of channels in regions of interest within the insular cortex. Notice that 8 channels were recording from two insular sub-areas and therefore the total number of channels (last column) might not be equal to the sum of the channels from the individual sub-areas (columns 4 to 8). SG: short insular gyri, LG: long insular gyrus and central sulcus of the insula, aCS: anterior segment of the circular sulcus of the insula, sCS: superior segment of the circular sulcus of the insula, iCS: inferior segment of the circular sulcus of the insula.

**Auditory effects: The responsive channels**

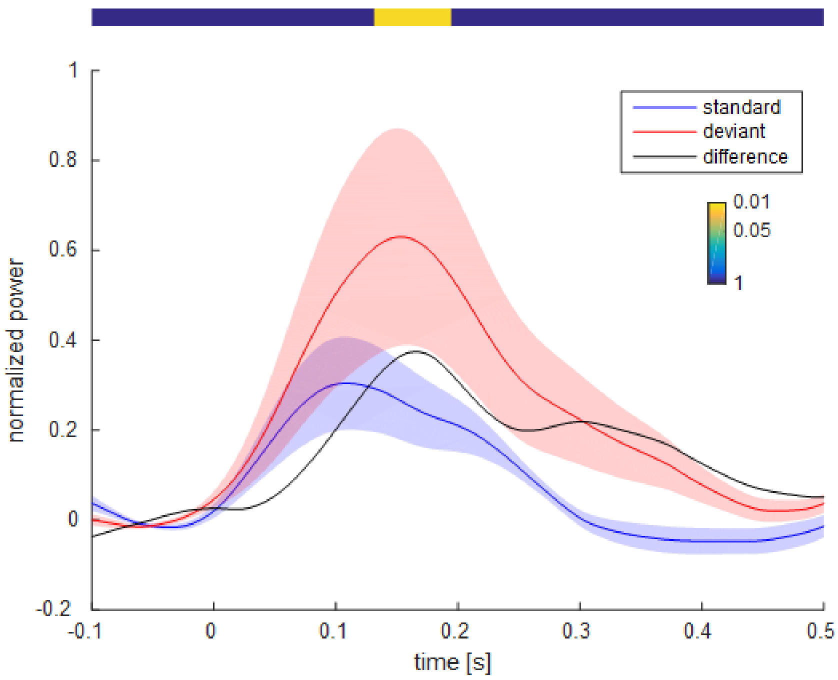


**Supplementary Figure 1**
Mean HFA response to standard and deviant tones in all Tone-responsive channels (n = 29). Black line shows the mean difference signal between deviants and standards. Response to deviants is higher between 134 and 196 ms after stimuli onset (p = 0.014, cluster-based permutation t-test). Shadowed areas depict mean +/- s.e.m.

**Deviant effects: The Condition-specific channels**

For further comparisons with other studies, we report the mean (x, y, z) coordinates in MNI space of the left and right Condition-specific channels to be located at (-42, -15, 5) and (42, -16, -3).

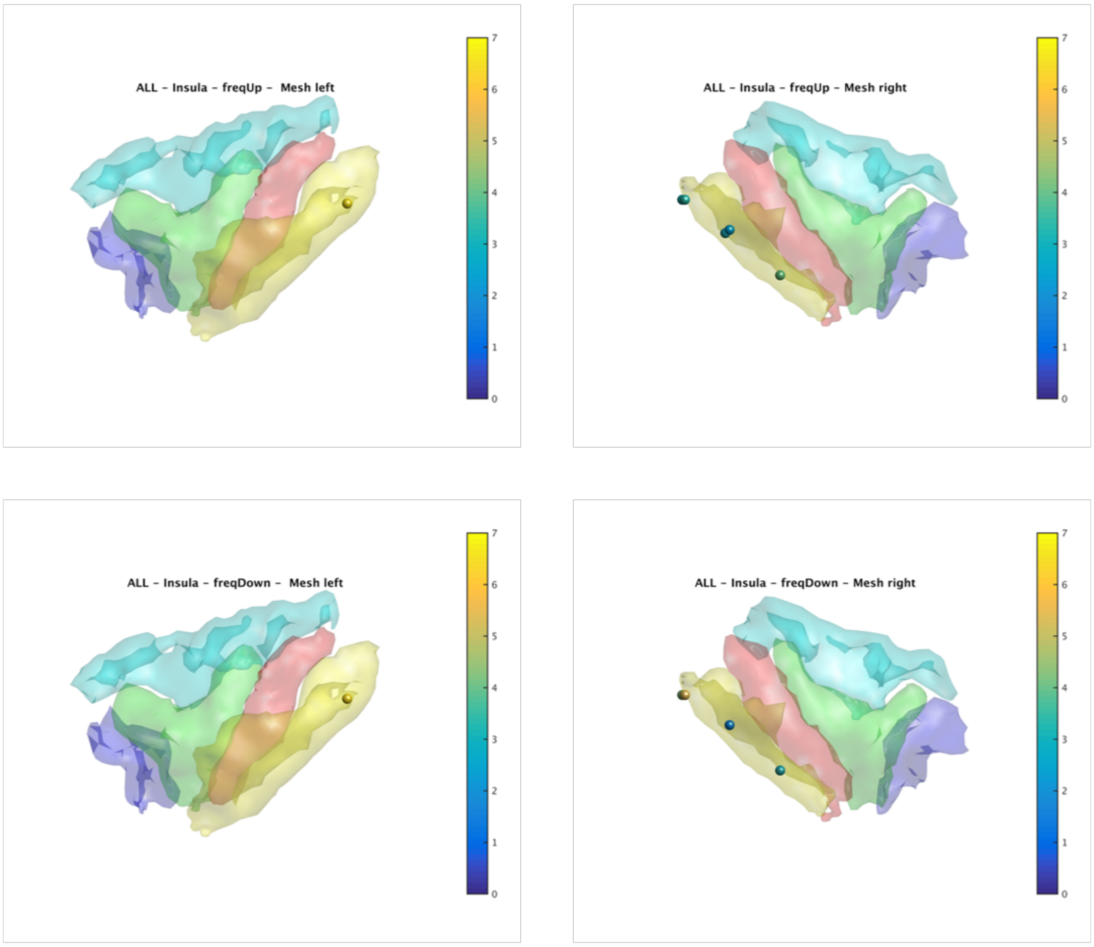


**Supplementary Figure 2**
Mean HFA responses to frequency up and down deviants (0-300 ms). Color bars represent t values. dof = 42 to 60, and dof = 49 to 57, respectively.

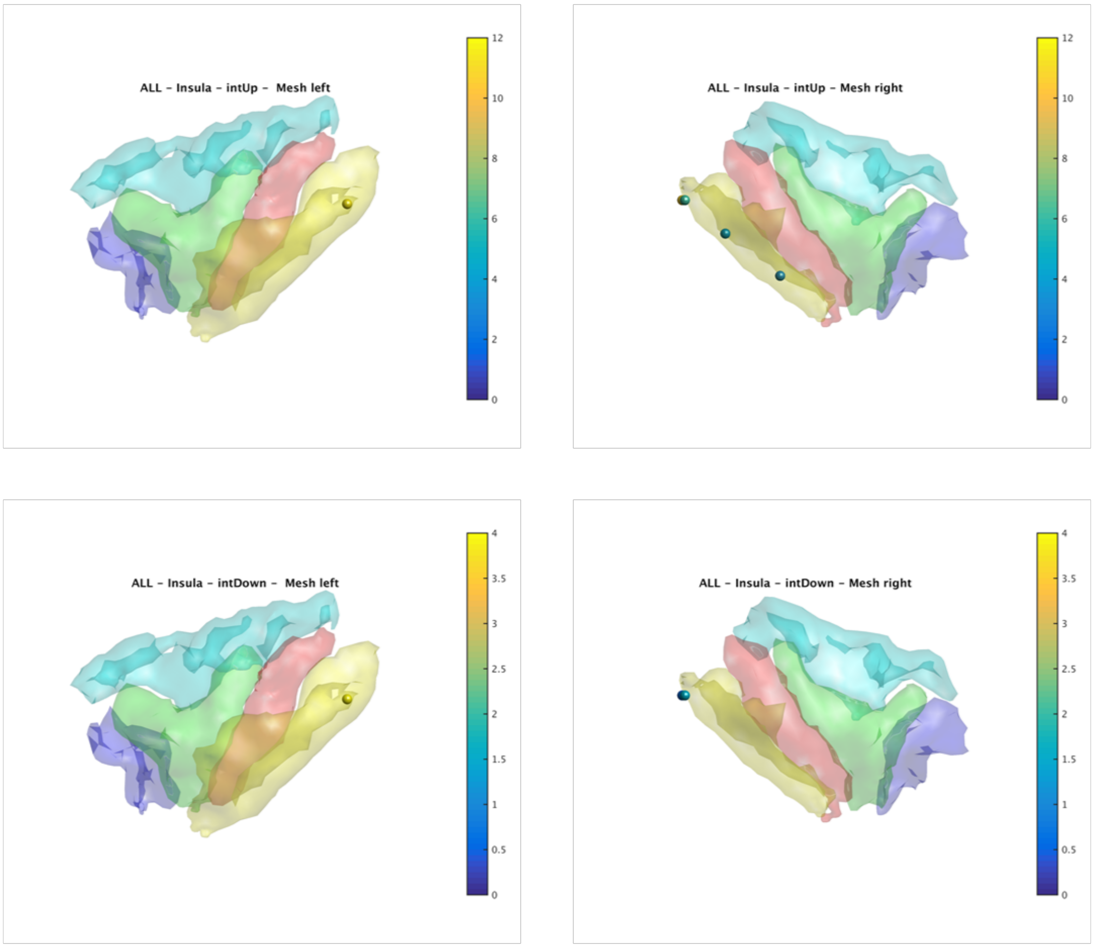


**Supplementary Figure 3**
Mean HFA responses to intensity up and down deviants (0-300 ms). Color bars represent t values. dof = 42 to 66, and dof = 60, respectively.

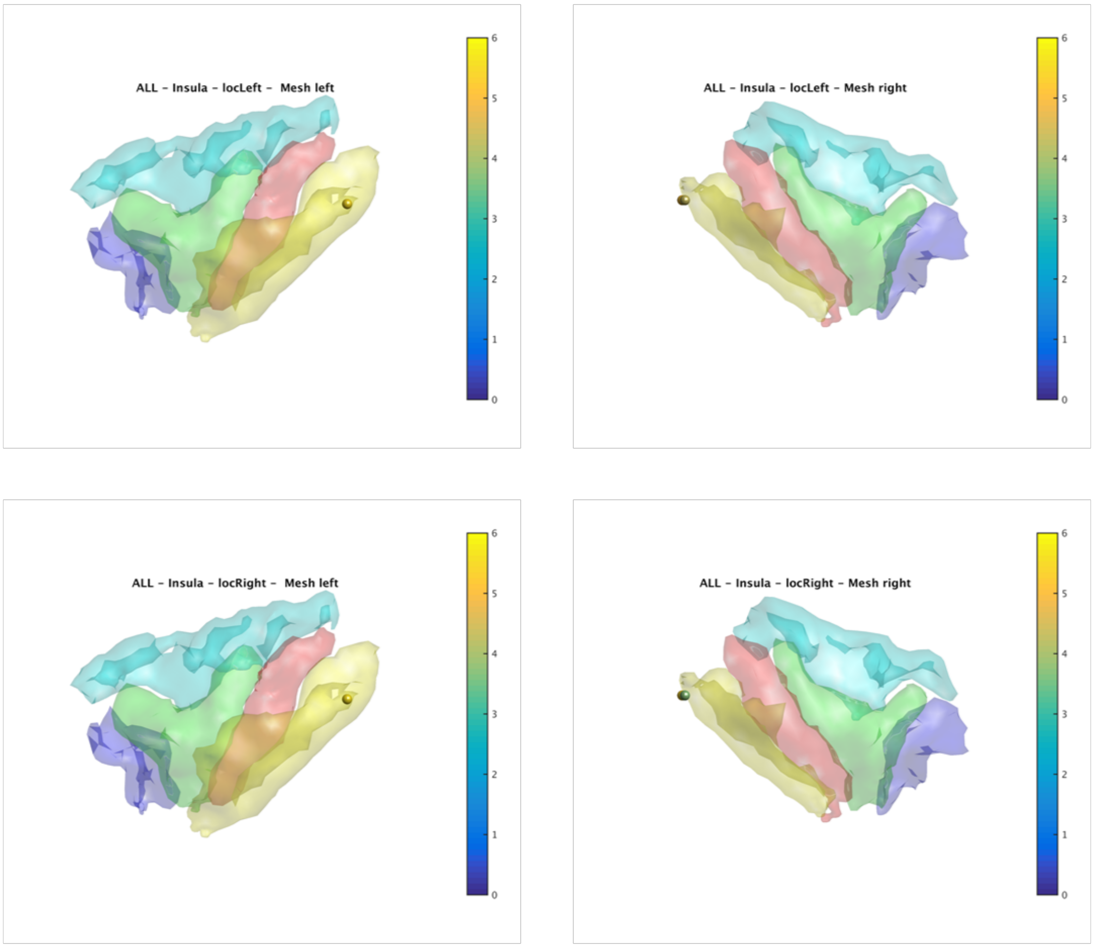


**Supplementary Figure 4**
Mean HFA responses to location right and left deviants (0-300 ms). Color bars represent t values. dof = 60 for both cases.

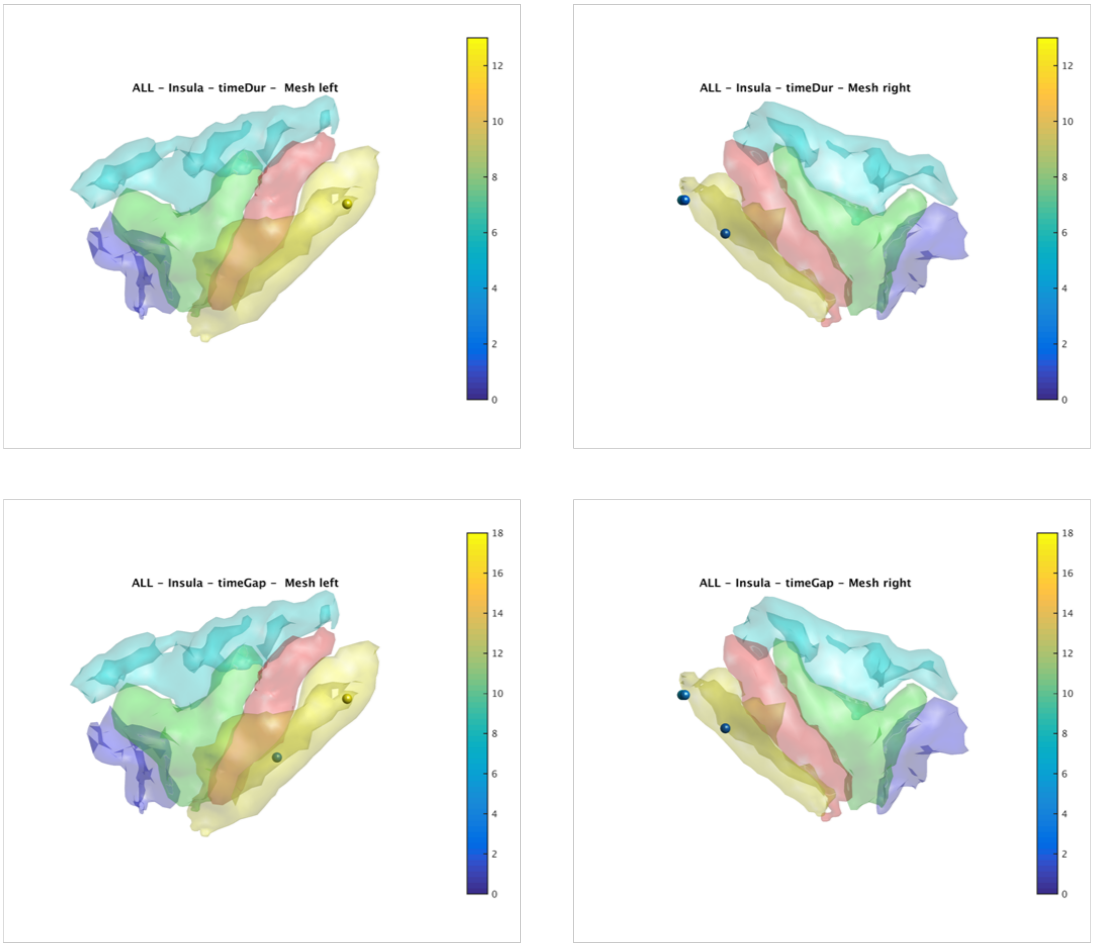


**Supplementary Figure 5**
Mean HFA responses to time duration and time gap deviants (0-300 ms). Color bars represent t values. dof = 83 to 119, and dof = 92 to 220, respectively.

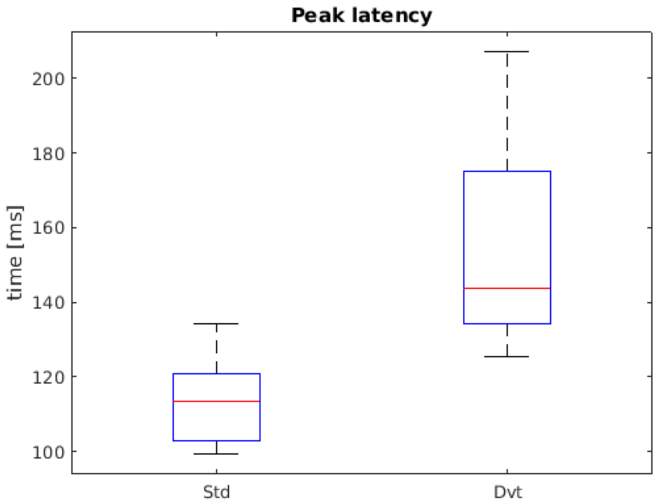


**Supplementary Figure 6**
Peak latency distribution of HFA signals in the Condition-specific channels to standard and deviant tones. Standard tones peaked earlier than deviants (median 105 vs. 143 ms, Wilcoxon signed rank test, $p = 0.008$).

**Post-Hoc analysis on the spatial distribution of responsive and condition specific channels in the inferior circular sulcus (iCS)**

All channels in the iCS were splatted in 3 groups of equal number of channels along the anterior-posterior *y* axis. The *anterior* group considered channels with *y* in the range [-3.35 6], the middle group with y in the range [-3.35 -9.33], and the posterior group with y in the range [-24 -9.33]. Coordinates are in MNI space.

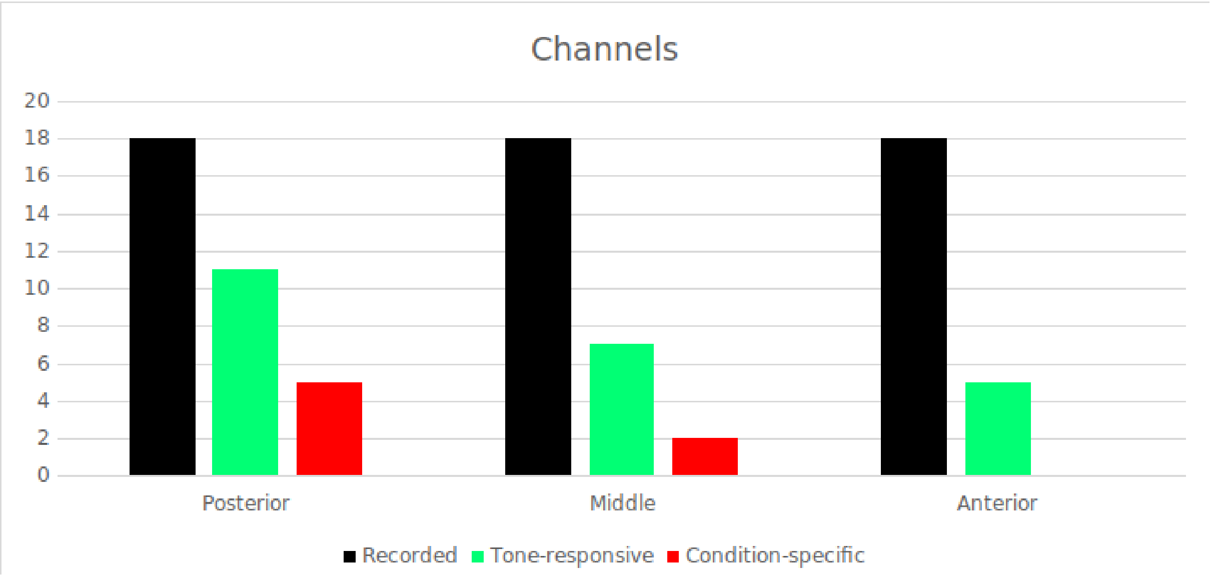


**Supplementary Figure 6**
Distribution of recording, responsive and condition specific channels across the anterior, middle and posterior part of the iCS.

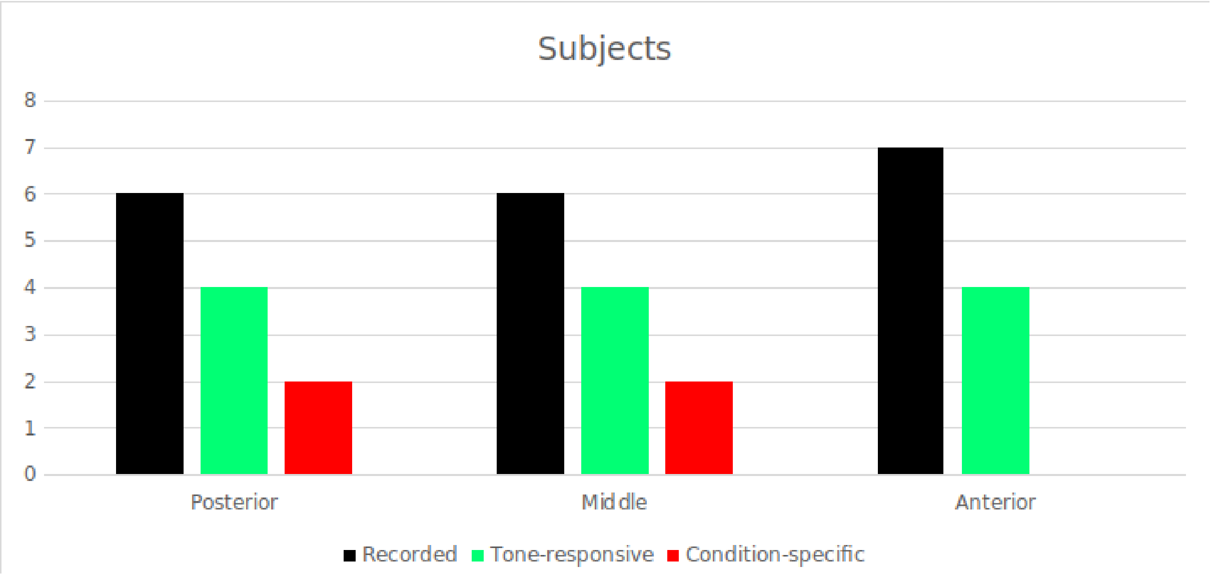


**supplementary Figure 7**
Distribution of subjects with recording, responsive and condition specific channels across the anterior, middle and posterior part of the iCS.
